# Computing, analyzing and comparing the radius of gyration and hydrodynamic radius in conformational ensembles of intrinsically disordered proteins

**DOI:** 10.1101/679373

**Authors:** Mustapha Carab Ahmed, Ramon Crehuet, Kresten Lindorff-Larsen

## Abstract

The level of compaction of an intrinsically disordered protein may affect both its physical and biological properties, and can be probed via different types of biophysical experiments. Small-angle X-ray scattering (SAXS) probe the radius of gyration (*R*_*g*_) whereas pulsed-field-gradient nuclear magnetic resonance (NMR) diffusion, fluorescence correlation spectroscopy and dynamic light scattering experiments can be used to determine the hydrodynamic radius (*R*_*h*_). Here we show how to calculate *R*_*g*_ and *R*_*h*_ from a computationally-generated conformational ensemble of an intrinsically disordered protein. We further describe how to use a Bayesian/Maximum Entropy procedure to integrate data from SAXS and NMR diffusion experiments, so as to derive conformational ensembles in agreement with those experiments.

## Introduction

In contrast to natively folded proteins, intrinsically disordered proteins (IDPs) generally lack well-defined three-dimensional structures. Consequently, they explore a large number of distinct conformations, and their conformational properties are thus best described in statistical terms. One useful and informative way of representing this large conformational ensemble is through a distribution of the radius of gyration (*R*_*g*_) of the IDP. The ensemble average 〈*R*_*g*_〉 gives a rough measure of how compact a protein is and may, for example, be compared to the values for other proteins of similar lengths.

For a given configuration of a protein, the *R*_*g*_ may be calculated as the mass-weighted root mean distance to the centre of mass:

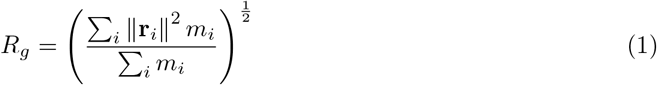

where *m*_*i*_ is the mass of atom *i* and **r**_*i*_ is the position of atom *i* with respect to the center of mass of the molecule.

Experimentally, one may obtain an estimate of the ensemble-averaged value of the *R*_*g*_ of a protein by a Guinier analysis of small angle X-ray scattering (SAXS) profiles (*1*) or using various extended models of the scattering data (*2*, *3*). For the sake of simplicity, we will loosely refer to the experimental value as *R*_*g*_, omitting the bracket notation and only use brackets for explicitly averaging computed values. Here, we note also that *R*_*g*_ calculated using Eq. 1 is not directly comparable to that obtained from analyses of SAXS data due to contributions to the scattering data from the solvent layer around the disordered protein (*4*, *5*).

Similarly, but via different physical principles, the hydrodynamic radius of a protein also reports on the overall expansion of a protein. The hydrodynamic radius (*R*_*h*_), also called the Stokes radius, is defined as the radius of a theoretical hard sphere that would have the same translational diffusion coefficient as the considered particle. The translational diffusion coefficient (*D*_t_) of a protein may in turn be determined e.g. by pulsed-field gradient Nuclear Magnetic Resonance (NMR) diffusion experiments, fluorescence correlation spectroscopy and dynamic light scattering measurements, and is related to *R*_*h*_ through the Stokes-Einstein equation (***6***):

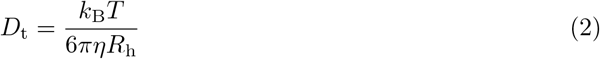

where *k*_B_ is the Boltzmann constant, *T* is the temperature and *η* is the viscosity of the solvent.

Because both *R*_*g*_ and *R*_*h*_ probe the compaction of a disordered protein, and because they may contain complementary information about the distribution of states (*7*) there have been several studies on the relationship between the *R*_*g*_ and *R*_*h*_ for disordered proteins and polymers (*7* –*10*).

One such approach uses hydrodynamic modelling of protein conformations (*11* –*13*) to relate protein structure to *R*_*h*_ (*7*, *10*). In line with theoretical expectations, the authors found that the ratio *R*_*g*_/*R*_*h*_ depends substantially on the compaction of the protein chain, so that compact states have ratios *≈* 0.8 and expanded conformations have ratios between 1.2–1.6. Because the relative level of compaction of the chain, when quantified by *R*_*g*_, also depends on the chain length, the ratio *R*_*g*_/*R*_*h*_ also depends on the number of residues of the protein (*N*). Recently, these two effects were combined into a single, physically-motivated and empirically parameterized equation that enables one to calculate *R*_*h*_ for a configuration of an IDP from its *R*_*g*_ (*14*):

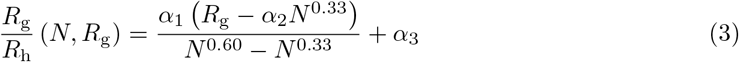

In addition to *R*_*g*_ and *N* (number of residues of the protein chain), the equation contains three parameters that were fitted to maximize agreement between the model and hydrodynamic calculations (*α*_1_ = (0.216 ± 0.001)Å, *α*_2_ = (4.06 ± 0.02)Å, and *α*_3_ = (0.821 ± 0.002)Å). As discussed further below, since conformational averaging acts on the diffusion properties, the ensemble averaged value that should be compared to an experimentally measured *R*_*h*_ will not in general be the same as the linear average over the values of each conformation (〈*R*_*h*_〉). Also, note that the equation was parameterized using *R*_*g*_ values calculated from the C_*α*_ coordinates only. Values of *R*_*g*_ calculated in this way are generally very close to those calculated from all protein atoms, but this parameterization makes it possible to use the approach to calculate *R*_*h*_ also for coarse-grained C_*α*_-only models.

Here we provide a step-by-step protocol to calculate *R*_*g*_ and subsequently *R*_*h*_ using Eq. 3 from a computationally generated conformational ensemble of an IDP. Together with calculations of SAXS data from simulations it is possible to compare the simulations to measurements of compaction. In cases where the computed and experimental quantities are not in perfect agreement, one may go one step further and refine the computational ensemble using the experimental data. We thus also demonstrate how to refine the ensembles by integrating experimental SAXS and *R*_*h*_ measurements, and thereby generate conformational ensembles that both take into account the physical principles encoded in the simulations as well as information from experiments. In addition to the motivation and description provided in this paper we also make available a Jupyter (Python) notebook with guided examples for performing analysis and generating many of the figures discussed here. We do, however, not provide instructions for how to generate conformational ensembles, and the reader is expected to have a basic understanding of the Python programming language to use the examples presented.

## 2 Materials

### Experimental data and sequence of Sic1

– We used the following sequence for the Sic1: GSMTPSTPPR SRGTRYLAQP SGNTSSSALM QGQKTPQKPS QNLVPVTPST TKSFKNAPLL APPNSNMGMT SPFNGLTSPQ RSPFPKSSVK RT
– SAXS data for Sic1 (*15*) obtained from the Protein Ensemble Database (*16*) entry PED9AAA. (http://pedb.vib.be/accession.php?ID=PED9AAA
– We used the previously measured (*17*) experimental value of *Rh* (21.5 *±* 1.1Å)

Software:

– Flexible Meccano (*18*) available from http://www.ibs.fr/research/scientific-output/software/flexible-meccano/?lang=en
– CAMPARI v3.0 (*19*) available from https://sourceforge.net/projects/campari/
– PULCHRA v3.06 (*20*) available from http://www.pirx.com/pulchra/index.shtml
– Pepsi-SAXS v1.4 (*21*) available from https://team.inria.fr/nano-d/software/pepsi-saxs/
– BME (*22*) available from https://github.com/KULL-Centre/BME
– MDtraj v1.9.3 (*23*) available from http://mdtraj.org/1.9.3/
– A Python Jupyter notebook (https://jupyter.org/) for performing the calculations and analyses described in this paper is available from https://github.com/KULL-Centre/papers/edit/master/2019/IDP-methods-Ahmed-et-al/

## 3 Methods

### 3.1 Generating Ensembles

We have chosen the 90 amino acid residues long protein Sic1 as an example for our calculations, as this protein has been studied extensively by both SAXS and various NMR methods (*15*, *17*). We used Campari (*19*) and Flexible-Meccano (*18*) to generate two conformational ensembles of Sic1 in its unphosphorylated state. In the ensemble we generated using Campari (Ensemble 1) we used Monte Carlo sampling with the ABSINTH v3.2 implicit solvent model (*24*) and a temperature of 298K. The Sic1 protein was contained in a spherical simulation cell with a radius of 150 Å and an ion concentration of ≈140 mM, matching the experimental condition (*15*). For the Flexible-Meccano ensemble we generated conformations sampling random coil configurations as described (*18*). As Flexible-Meccano only generates a model of the protein backbone, we used PULCHRA (*20*) with default settings to add side chains to these structures and generate Ensemble 2. These side chain coordinates are necessary when we calculate SAXS data from the conformational ensembles. In total we generated 32,000 structures for Ensemble 1 and 10,000 structures for Ensemble 2.

### 3.2 Calculating Rg and Rh from ensembles

Many simulation and protein analysis software packages have the option of calculating the *R*_*g*_ of the protein. In this example we will use readily available and open source software. For calculating the *R*_*g*_ of the conformations we use MDTraj, a python module for protein analysis (*23*). Below we provide Python code demonstrating how to load the ensemble and calculate *R*_*g*_ for each structure, and then calculate *R*_*h*_ for each structure using Eq. 3. In the example we have collected all conformations of the ensemble in a trajectory file (here Ensemble1.trr). Depending on the file format of the trajectory file, one may also need a coordinate file (structure.pdb) or a topology file. Once these files are loaded, MDTraj is then used to calculate *R*_*g*_ for each structure in the ensemble, which in turn is converted into *R*_*h*_ using Eq. 3.

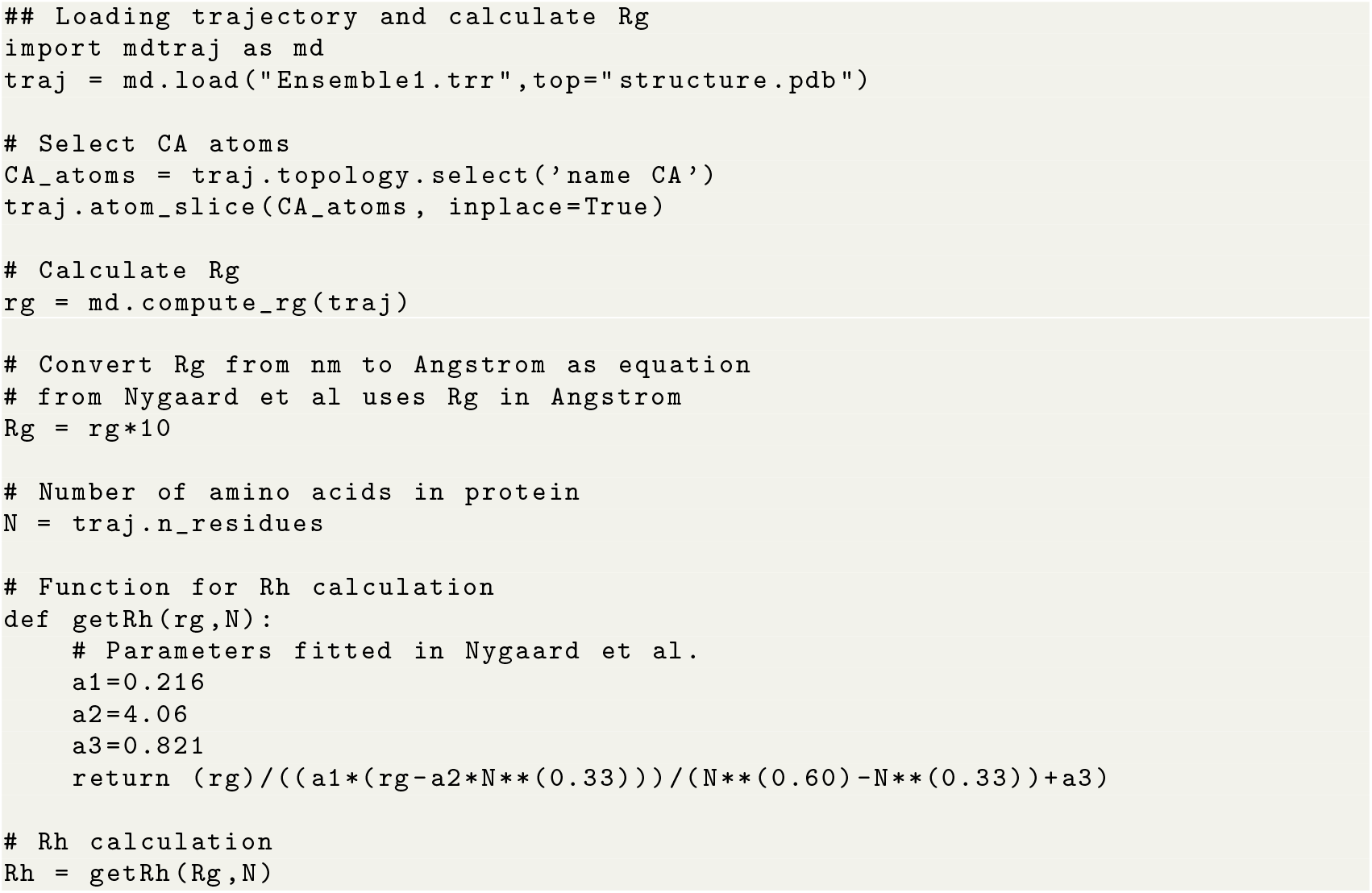

Once *R*_*g*_ and *R*_*h*_ have been calculated for each structure, these can be used to generate histograms of *R*_*g*_ (Fig. 1a) and *R*_*h*_ (Fig. 1b), and the average *R*_*h*_ can be calculated as for comparison to experimental values (see Note 1). We also show the calculated average of *R*_*g*_ in the plots (Fig. 1) (see Note 2) though as explained below, a better comparison to the experimental data requires calculations of SAXS intensities from the conformational ensemble (see also Note 3).

**Fig. 1.**
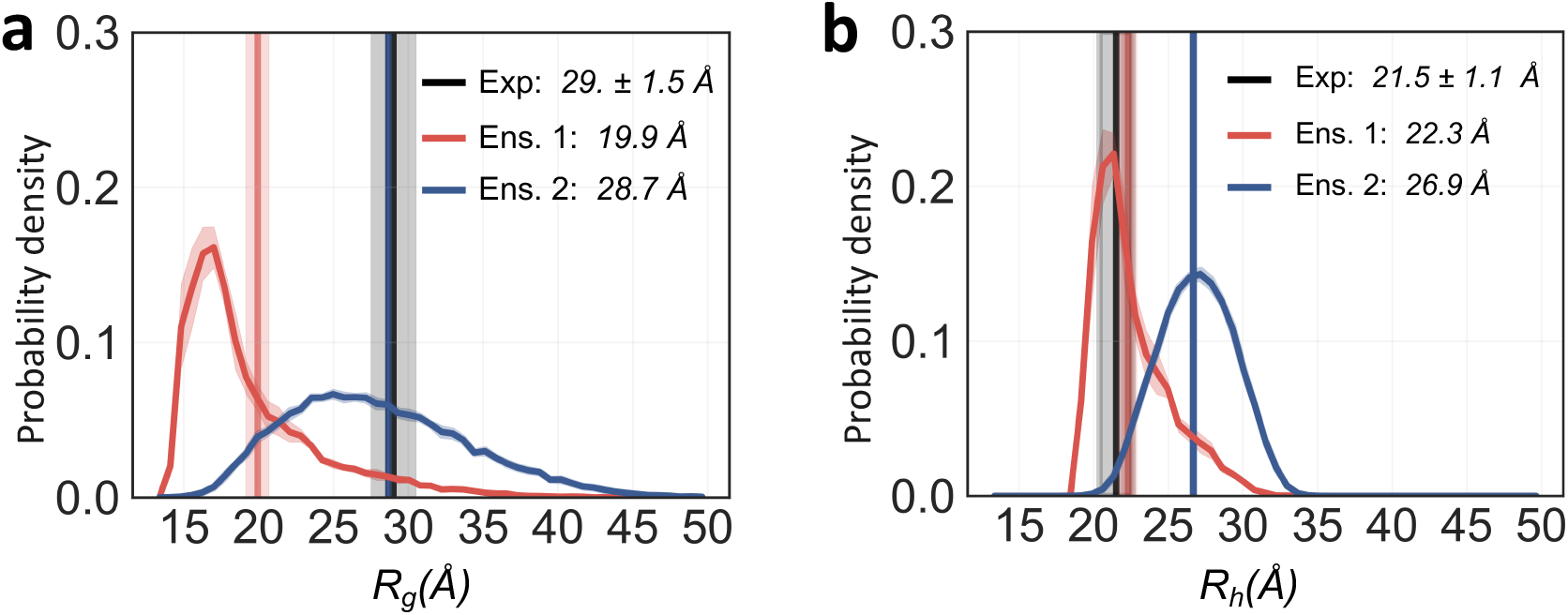
Analyzing compaction in ensembles of Sic1. Probability distribution of **(a)** *R*_*g*_ and **(b)** *R*_*h*_ calculated from two ensembles that we generated of Sic1. Here, Ensemble 1 (red) was generated using Campari and Ensemble 2 (blue) was generated using Flexible-Meccano as described in the main text. **(a)** Solid vertical lines represents the ensemble average *R*_*g*_ (〈*R*_*g*_〉_*trans*_; see Note 2 for the definition) of Ensemble 1 (red) and Ensemble 2 (blue). **(b)** Solid vertical lines represents the ensemble average *R*_*h*_ (〈*R*_*h*_〉_*trans*_) calculated using Eq. 3 and as discussed in Note 1 from Ensemble 1 (red) and Ensemble 2(blue). The experimental values of *R*_*g*_ and *R*_*h*_ are shown in black. The error of the distribution and averages of *R*_*g*_ and *R*_*h*_ (shown as shaded areas) were estimated by block averaging using five blocks.

As described above the ratio *R*_*g*_/*R*_*h*_ depends substantially on the compaction of the protein chain, so that compact states have ratios *≈* 0.8 and expanded conformations have ratios between 1.2–1.6 (*14*). For a protein of 91 amino acids, the switch-over point where the *R*_*g*_/*R*_*h*_ = 1 lies at conformations with *R*_*g*_ ≈ 27Å (see Note 4). Thus, conformations with *R*_*g*_ < 27Å have *R*_*h*_ > *R*_*g*_ whereas conformations with *R*_*g*_ > 27Å have *R*_*h*_ < *R*_*g*_. In this way, the distribution of *R*_*h*_ is ‘pushed’ towards the middle and has less density in the tails compared to the distribution of *R*_*g*_ (Fig. 1).

The distributions of *R*_*g*_ (Fig. 1a) and *R*_*h*_ (Fig. 1b) from Ensemble 1 and Ensemble 2 and the resulting averages, can also be compared to the experimental values from SAXS and NMR (*15*, *17*). These results reveal two different scenarios for the two ensembles. First, 〈*R*_*h*_〉_*trans*_ calculated from Ensemble 1 (Campari) is in good agreement with the experimentally-determined value of *R*_*h*_ (Fig. 1b). At the same time, the calculated value of 〈*R*_*g*_〉_*trans*_ is substantially lower than the average *R*_*g*_ value estimated from SAXS experiments (Fig. 1a). Second, for Ensemble 2 (Flexible-Meccano) we observe the opposite scenario, where the calculated 〈*R*_*g*_〉_*trans*_ is close to the value estimated by SAXS (Fig. 1a), and the calculated 〈*R*_*h*_〉_*trans*_ is substantially greater than the experimental value (Fig. 1b).

Disagreement between experiment and simulation is often indicative of problems with the molecular force fields or sampling (*25*), though differences may also arise from problems in e.g. the model used to calculate experimental data from structural ensembles (*5*, *26*). While it is possible to improve molecular force fields directly against experimental data (*27*), we below describe how one can refine a specific ensemble against one or more sets of experimental measurements.

### 3.3 A Bayesian/Maximum Entropy approach

Above we have analysed two ensembles and used Eq. 3 to estimate *R*_*h*_ which in turn could be averaged and compared to NMR diffusion experiments. We also calculated *R*_*g*_ from the protein co-ordinates, though as noted this value is not directly comparable to the experimental measurements due to solvation effects (*5*). Nevertheless, the results suggested discrepancies between experiments and simulations.

Although there has been continued improvements in methods and force fields for sampling the conformational landscape of IDPs, it is still not uncommon that simulations are not in perfect agreement with experiments. In such cases, it is possible to bias the simulation to construct an ensemble that is in better agreement than the unbiased ensemble (*22*, *28* –*31*).

We here use such a method to construct two new ensembles by reweighting the Campari and Flexible-Meccano ensembles with the experimental data, thus obtaining ensembles that are in better overall agreement with the SAXS and NMR diffusion experiments. Specifically, we use experimental SAXS data (*15*) and NMR diffusion measurements of *R*_*h*_ (*17*), and use our recently described Bayesian/Maximum Entropy (BME) protocol to reweight the conformational ensembles (*22*). We focus solely on the technical details of the approach rather than the biological relevance. Also, we exemplify using two experimental measures of compaction, but the approach is more generally applicable (See Note 5).

Briefly described, BME is based on a combined Bayesian/Maximum Entropy framework, and enables one to refine a simulation using multiple sources of (potentially noisy) data. The purpose of the reweigthing is to derive a new set of weights for each configuration in a previously generated ensemble so that the reweighted ensemble satisfies two criteria: (i) it matches the experimental data better than the original ensemble and (ii) it achieves this improved agreement by a minimal perturbation of the original ensemble. For additional details see Bottaro et al. (*22*) and references therein. In the current examples, both ensemble 1 and 2 were generated as unbiased ensemble and so the initial weights of all structures are uniform 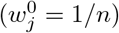, where *n* is the number of structures in the ensemble.

The reweighting approach described above may in practice be achieved by updating the weights, *w*_*j*_, of each configuration in the input ensemble by minimizing a function (the negative log-likelihood) (*22*, *28*):

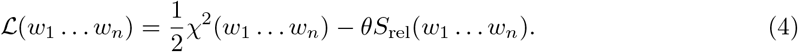

Here, the *χ*^2^ quantifies the agreement between the experimental data and the corresponding values calculated from the reweighted ensemble. The second term contains the relative entropy, *S*_rel_, which measures the deviation between the original ensemble and the reweighted ensemble 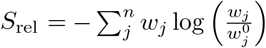. The temperature-like parameter *θ* tunes the balance between fitting the data accurately (low *χ*^2^) and not deviating too much from the prior (low *S*_rel_). In practice, we determine this hyperparameter by evaluating the compromise between balancing the two terms in 𝓛 (*22*, *28*) (see also Note 6). When more than one set of experimental data is included in BME, the deviations between calculated and experimental values are summed in a global *χ*^2^ function which is the sum of a *χ*^2^ function for each set of data.

In practice it turns out that in many cases there is a more efficient approach to minimize 𝓛 using the method of Lagrange multipliers, and this is the approach we take here (*22*, *28*, *32*) using the BME code, which is freely available at https://github.com/KULL-Centre/BME.

### 3.4 Calculating SAXS data from ensembles

The first step in the reweighting protocol is to collect the necessary data and structure it correctly for input in BME. We first calculate the SAXS intensity profiles by fitting to the experimental curve for each structure of the two ensembles using Pepsi-SAXS (*21*). Pepsi-SAXS has free parameters for the solvation layer that are calculated for each fit. To decrease the risk of overfitting, we used a two-step procedure. First, we fitted the parameters to each structure. Second, we calculate the averages of the resulting fitted values of the solvation parameters and re-ran Pepsi-SAXS with these parameters fixed to those averages. Alternative methods for calculating SAXS from conformational ensembles exist (*4*) and may also be used (See Note 7 and Note 8).

We then structure the input files as shown below for SAXS BME input. The experimental SAXS input file is structured such that it contains the following three columns: the momentum transfer (*q*), intensity (*I*(*q*)), and the error (*σ*_*I*_ (*q*)) (as shown below). Each of these three columns are *m* rows long, where *m* is the number of experimental data points. The input file for the calculated values contains *n* rows(number of structure in the ensemble), and *m* + 1 columns. The first column is for labeling the individual structure/frame from the ensemble. Further details for how to structure the input files for other data can be found in the original description of BME and in the online examples (*22*).

#### Experimental file format

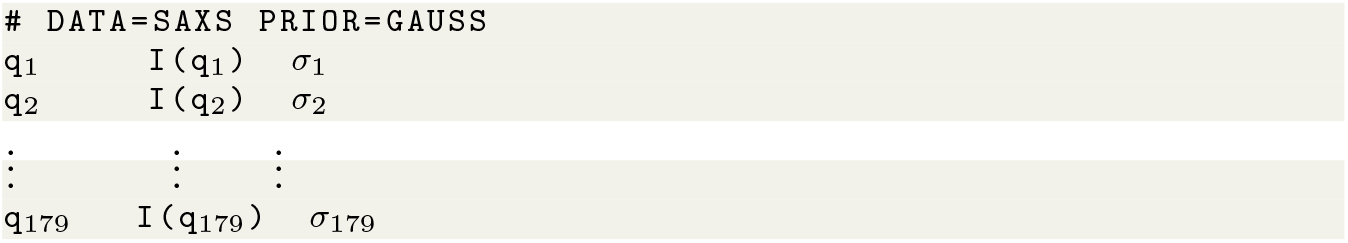

#### Simulation SAXS file format

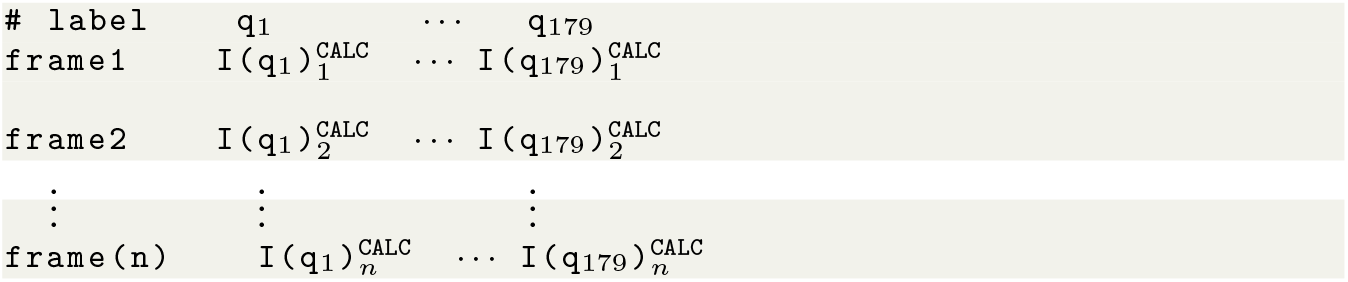

Once these calculations have been done, we may load the data in python and run BME:

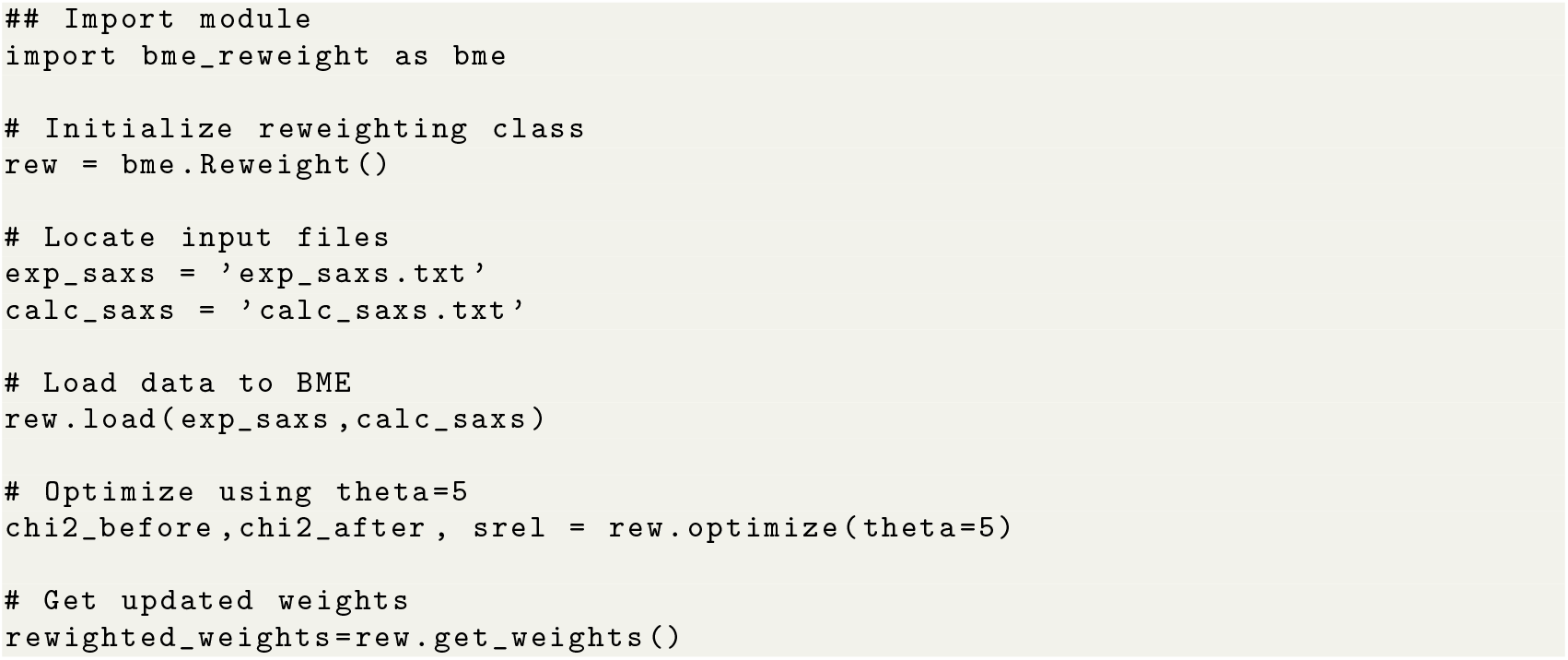

### 3.5 Reweighting Sic1 ensembles against SAXS and NMR diffusion experiments

We used the methods described above to determine a reweighted ensemble of Sic1 that takes into account both the prior information encoded in the initial ensemble (from Campari or Flexible-Meccano) as well as the experimental measurements of compaction from NMR diffusion and SAXS.

Before reweighting was applied, Ensemble 1 appears too compact when judged by agreement with the *R*_*g*_-value extracted from the SAXS data, but is in good agreement with the NMR diffusion data (Fig. 1). In contrast, Ensemble 2 is in good agreement with the SAXS-derived *R*_*g*_, but appears too expanded when compared to the NMR diffusion measurements (Fig. 1). The goal was therefore to examine whether one could construct an ensemble that provides a useful compromise between the two data sets. We note here that the NMR diffusion data were recorded at 278 K (*17*), whereas the SAXS data were obtained at room temperature (*15*), though we only expect a modest change in compaction in this temperature range (*33*, *34*). We note also that our goal is not to discuss in detail the conformational ensemble of Sic1, but rather to showcase how one may combine different measures of compaction.

We reweighted the two ensembles against the NMR and SAXS data and compared to the unweighted ensembles (Fig. 2). The first step is to chose the temperature-like hyperparameter, *θ*, that sets the balance between fitting the data and not deviating too much from the input ensemble. The latter may be quantified by calculating the fraction of the frames in the input ensemble, *N*_*eff*_ = exp(*S*_*rel*_), that effectively contribute to the calculated ensemble averages after reweighting. Thus, *N*_*eff*_ = 1 corresponds to the initial unweighted ensemble and a low value of *N*_*eff*_ indicates that only a small fraction of the original ensemble has been selected to improve agreement with experiments. We scanned values of *θ* and calculated the agreement with both the SAXS and NMR diffusion data at each value of *θ* and for each of the two ensembles (Figs. 2a and 2b). Note that we here plot a reduced 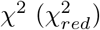 for each of the two experiments individually, but that the optimization acts to reduce the sum of the two non-reduced *χ*^2^-values. Since there is a 179 points in the SAXS measurements, this sum contains a large contribution from the SAXS data (see Note 8). In our analyses here, we chose *θ* = 100 for Ensemble 1 and *θ* = 7 for Ensemble 2, though in practical applications it would be advised to examine the results of other choices (See Note 6).

**Fig. 2.**
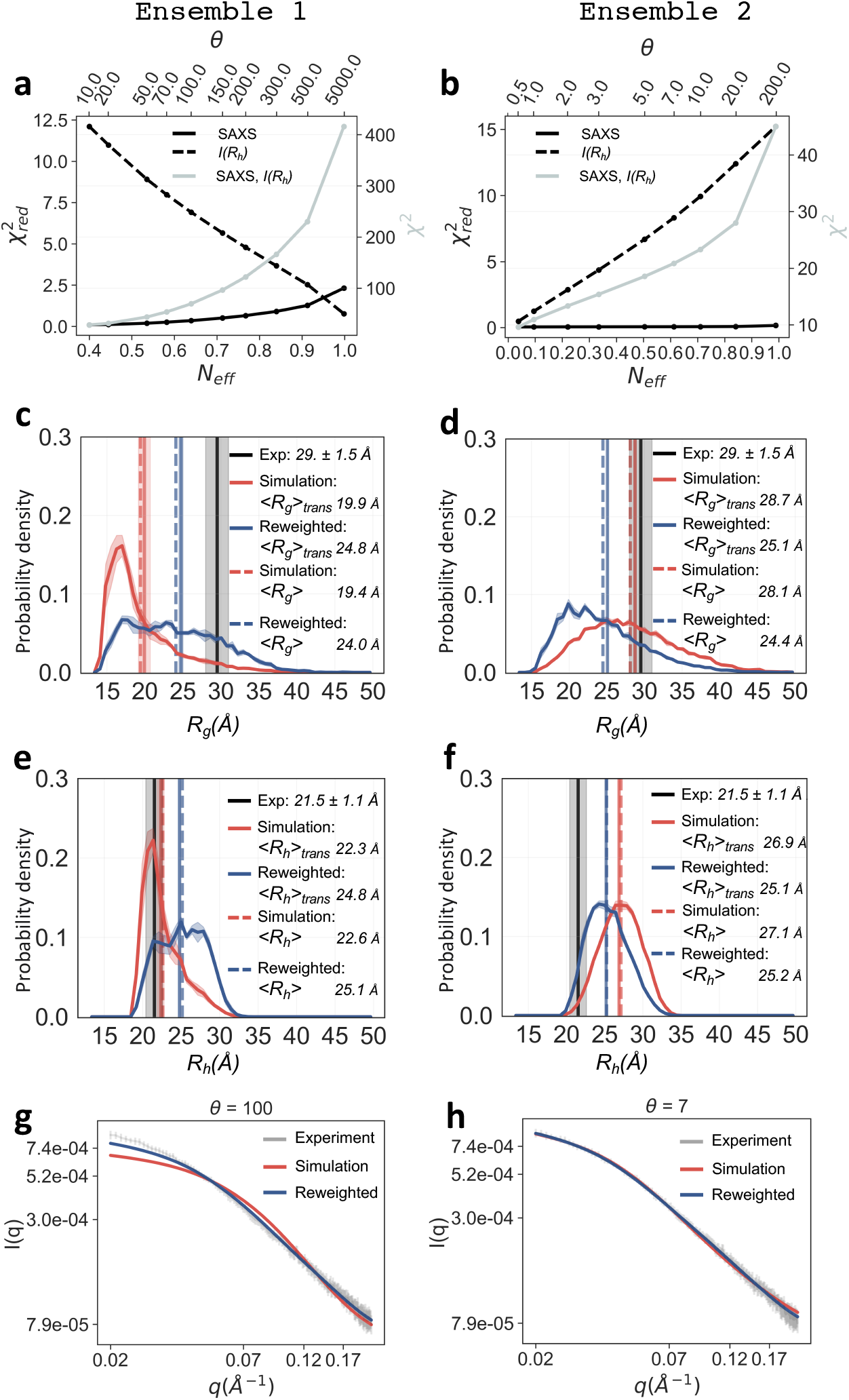
Constructing ensembles to improve agreement with experiments. We used BME reweighting with SAXS and *R*_*h*_ data for Ensemble 1 (**a, c, e, g**) and Ensemble 2 (**b, d, f, h**). We label the *R*_*h*_ data as *I*(*R*_*h*_) as we here use intensity-based averaging of the measurements (Note 1). (**a,b**) We plot *N*_*eff*_ (the effective number of frames left after reweighting) vs. *χ*^2^ when the scaling parameter *θ* is varied (top axis). The left axes show 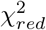 for each of the two experiments, whereas the right axis shows the total *χ*^2^ that is the sum of the two (non-reduced) *χ*^2^ values (Note 8). For further analyses we chose *θ* = 100 (Ensemble 1) and *θ* = 7 (Ensemble 2). (**c-f**) We show the distribution of *R*_*g*_ (**c,d**) and *R*_*h*_ (**e,f**) before (red) and after (blue) reweighting the two ensembles. Averages over these distributions (both before and after reweighting) are shown either as standard (linear) averages (dashed lines) or ‘transformed’ averages (〈*R*_*g*_〉_*trans*_ and 〈*R*_*h*_〉_*trans*_ as described in Notes 1 and 2). (**g,h**) We show the calculated SAXS intensity from the original ensemble and the refined ensembles and compared to the experimental data. In panels c–f the experimental data are shown in black lines and the errors are shown as shades. Errors in calculated values were estimated by block averaging using 5 blocks.

The effect of reweighting can be seen both on the distribution of *R*_*g*_ (Fig. 2c and 2d) and *R*_*h*_ (Fig. 2e and 2f). The more compact Ensemble 1 is shifted to include more expanded structures, bringing 〈*R*_*g*_〉_*trans*_ substantially closer to the value estimated from the SAXS data, while only increasing the calculated *R*_*h*_ value ≈ 15% above the experimental value. Similarly, the more expanded Ensemble 2 is shifted to give greater weight to more compact configurations, bringing the calculated *R*_*h*_ closer to experiment while only shifting the 〈*R*_*g*_〉_*trans*_ down by *≈* 13%.

While it is convenient to examine the distribution of *R*_*g*_ before and after reweighting, the actual reweighting is done against the SAXS data not the estimated *R*_*g*_. As explained above, the solvent layer around the protein also contributes to the SAXS measurements, and there may be *≈*5–10% difference in the *R*_*g*_ calculated from the protein coordinates and the value estimated by SAXS (*5*). We thus also show the agreement between the experimental and calculated SAXS curves (Fig. 2g and 2h). It is clear that the reweighted SAXS curves are substantially closer to the experimental data, though there still remains some discrepancy in the low-*q* range for Ensemble 1.

### 3.6 Summary

We have shown here how it is possible to calculate *R*_*h*_ from a conformational ensemble using Eq. 3 and compare to experimental data obtained e.g. from NMR diffusion measurements. Such measurements provide an alternative view of the compaction to that obtained e.g. from SAXS experiments, and indeed it has previously been shown that simultaneous refinement against *R*_*h*_ and *R*_*g*_ can provide insight into the shape of the distribution of *R*_*g*_ (*7*).

We chose the protein Sic1 for exemplifying our analyses since the level of expansion has been measured for this protein using both SAXS and NMR diffusion measurements. Since the data were recorded at slightly different conditions and temperatures, we do not aim to make strong conclusions about the conformational ensemble of Sic1, and have used it here mostly to showcase the methods for analyses.

We generated two ensembles and show that one is in relatively good agreement with the NMR diffusion data whereas the other is in better agreement with the SAXS data. At this moment the origins of these differences are unclear. Variation in experimental conditions such as temperature may affect both *R*_*g*_ and *R*_*h*_ (*34*). Also, it is possible that our approach for calculating *R*_*h*_ is not always sufficiently accurate since it is inherently limited to the accuracy achievable by hydrodynamic modelling (*14*), and an important question for future research is whether we can provide better models to link conformation and calculated values of *R*_*h*_. Finally, despite continued improvement in methods for calculating SAXS data from ensembles (*4*) there are still potential sources of error from e.g. solvation effects (*5*). Nevertheless, we note that by reweighting the ensembles against both sets of experiments it is possible to construct an ensemble that provides a reasonable balance between the two. As more proteins are studied by both NMR and SAXS it should be possible to test and improve our relationship between *R*_*g*_ and *R*_*h*_, thus enabling further insight into the rules that govern compaction of IDPs.

## 4 Notes

1. When calculating averages over ensembles, in particular for broad ensembles such as for IDPs, it is important to take the correct form of averaging into account. The best way to calculate averages over experimental quantities will depend both on the type of experiment and often also e.g. on the time scales for conformational averaging. Throughout this paper we make the assumption that averages can be calculated as time-independent averages over the conformational ensemble. In the case of measurements of the hydrodynamic radius, *R*_*h*_, we have explored two different types of averaging. In case the experiment measures the average diffusion coefficient, then according to Eq. 2 then the average should be calculated as 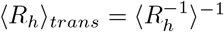. Here we 〈*R*_*h*_〉_*trans*_ have introduced the notation to represent that the averaging takes place on a *transformed* value (in this case proportional to 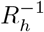). When *R*_*h*_ is measured by pulsed-field gradient NMR diffusion measurements (***35***) the NMR signal intensity, *I*, is proportional to 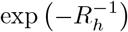 and it may therefore be more appropriate to use this function to perform the averaging. In this case

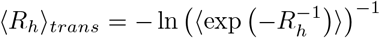 It is this intensity-based averaging that we use here, though in practice we have found it to give essentially the same result as using 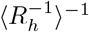.
2. Similar to the issue of averaging *R*_*h*_ discussed in Note 1 above, we use 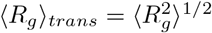 when calculating averages over the radius of gyration. This kind of averaging mimics the averaging in the low-*q* range of SAXS curves. Note, however, that 〈*R*_*g*_〉_*trans*_ calculated in this way should not directly be compared to experimental values of *R*_*g*_ since the latter includes solvation effects.
3. Notes 1 and 2 discuss the transformations that are relevant for comparing calculated and experimental quantities. We note, however, that during the reweighting protocol and generally when one makes quantitative comparisons between experiments and computation it is in general better to compare to the direct experimental quantities. In the case of SAXS experiments we thus judge agreement and perform reweighting against the experimentally measured intensities. In the case of the *R*_*h*_ measured for Sic1 by NMR diffusion experiments, we transform the experimental value of *R*_*h*_ (and its error), as well as the values calculated for each structure using the function 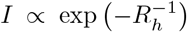, as described in our associated Jupyter notebook. We note that in the future it might be more appropriate to perform such fitting to the measured intensities as a function of the gradient strength.
4. The level of compaction as quantified by the value of *R*_*g*_ at which 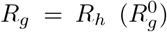 can be estimated by rearranging Eq. 3 to obtain: 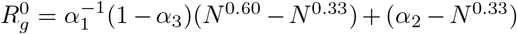. For a protein with *N* = 91 one obtains 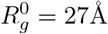.
5. We have here described approaches to refine ensembles against SAXS and NMR diffusion measurements. The BME method has also been used for IDPs with NMR chemical shifts (*36*), and may also readily be applied to SANS data, NOEs, scalar couplings or other measuremenets that can be calculated as averages over configurational ensembles.
6. Currently, the value of the hyperparameter *θ* (which sets the balance between information from the data and the force field) is set manually. In certain cases it may be possible to set it via a cross-validation approach (*36*) or may be integrated out as a Bayesian ‘nuissance parameter’ (*28*)
7. We have here used Pepsi-SAXS to calculate X-ray scattering curves from a conformational ensemble due to its ease of use and the relatively high computational efficiency. The latter is particularly important for large conformational ensembles. We note, however, that several other methods exist and suggest users in particular to keep solvent effects in mind when calculating and interpreting SAXS data (*4*, *5*). In the Jupyter notebook avialable online we provide a script that performs a two-pass run of Pepsi-SAXS to find a reasonable value of solvent-related parameters in the calculations.
8. When plotting 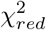 in Fig. 2, we calculate it by normalizing *χ*^2^ by the number of experimental data points: 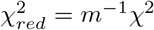. We note that this an approximation because the number of degrees of freedom can be smaller because different parameters are fitted such as parameters involved in calculated the SAXS curves. Also, in the case of reweighting the weights themselves may be considered as free parameters. Thus, we note that the reweighting does not involve this normalization, and that the 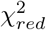 is only shown in Fig. 2 to give the reader an impression of the level of agreement. We also note that when fitting the *R*_*h*_ the resulting sum in *χ*^2^ only contains a single term. Finally, we note that we here simply combine the *χ*^2^ from the SAXS and NMR diffusion experiments by adding up the two individual *χ*^2^ terms. In the current implementation, BME does not enable automatic balancing of independent experiments and instead sets this balance by the error estimates of the individual experiments. We note, however, that while the SAXS data for Sic1 contains 179 individual data points, the amount of information in a SAXS experiments typically corresponds to a smaller number of parameters (*37*) and a more careful balance between the information in the SAXS and NMR diffusion experiments should take such effects into account (*38*).

## 5 Acknowledgements

We thank Dr. Tanja Mittag for providing feedback on the manuscript, Dr. Andreas Haahr Larsen for general discussions about SAXS experiments and calculations, and Dr. Martin Blackledge for suggesting to use intensity-based averaging for *R*_*h*_. The research described here was supported by a grant from the Lundbeck Foundation to the BRAINSTRUC structural biology initiative.

